# Development of a liquid biopsy for bladder cancer using a mutant protein panel in urinary extracellular vesicles

**DOI:** 10.1101/2025.05.12.653366

**Authors:** Yuji Hakozaki, Kazuma Sugimoto, Yuta Yamada, Tetsuya Danno, Kenichi Hashimoto, Hiroki Shinchi, Haruki Kume, Koji Ueda

## Abstract

Non-muscle invasive bladder cancer often relapses after cystoscopic surgery, necessitating rigorous monitoring for recurrence through invasive and painful cystoscopy. To develop a novel non-invasive and cancer-specific diagnostic method, somatic mutant proteins in urinary extracellular vesicles (EVs) were for the first time investigated using a proteogenomics pipeline consisting of whole exome sequencing and LC/MS. The analysis of bladder cancer tissues, cultured tissue-derived EVs, and urinary EVs from five patients identified 11,207, 9,809, and 5,828 unique proteins, respectively. Notably, 39, 32, and 4 mutant proteins were found in each of the sample sets. Furthermore, mass spectrometric absolute quantification measurements were conducted using prospectively collected urine samples, revealing that the levels of all monitored mutant proteins (LCP1_D321H, TKT_K102N, and PLCD1_R639H) exhibited a clear correlation with the cystoscopic tumor burden. Therefore, the presence of mutant proteins in Evs presents an ideal approach for liquid biopsy, serving as a non-invasive urine test for bladder cancer.

**Teaser:** Mutant proteins in urine were identified using a proteogenomic pipeline, which may serve as a novel method for monitoring bladder cancer.

## Introduction

Bladder cancer accounts for 3% of global newly diagnosed cancers, according to GLOBOCAN data (*1*). The disease is more common in men compared to women, and the 9^th^ leading cause of cancer death among men (*1*). Bladder cancer consists of two entities: non-muscular invasive (NMIBC) and muscle-invasive bladder cancer (MIBC). While MIBC is treated by cystectomy with neo-adjuvant chemotherapy, NMIBC can be managed with minimally invasive transurethral resection of the tumors along with the administration of Bacillus Calmette-Guerin or mitomycin C. However, intravesical recurrence will occur with these treatment strategies after surgery, and intensive monitoring is required to detect recurrence in the early stage. Based on the Surveillance, Epidemiology, and End Results (SEER)-Medicare database, the 10-year recurrence rate was 74.3% with bladder cancer patients, and it has the highest recurrence rate within solid tumors (*2*).

Cystoscopy and urine cytology are the standard methods to evaluate recurrence. National Comprehensive Cancer Network guidelines recommend cystoscopy and urine cytology checks every three months within two years after treatment for high-risk tumors. However, these surveillance methods are limited by their sensitivity, specificity, invasiveness, and care costs. Urine cytology is a non-invasive biomarker to detect recurrence of NMIBC. While the specificity is reported as 86%, the sensitivity was 48% to detect recurrence (*3*). To improve the sensitivity of the cytology check, cytoscopic follow-up is recommended for follow-up of patients. The sensitivity and specificity of cystoscopy were reported as 87% to 100% and 64% to 100%, respectively. Though the diagnostic performance of cystoscopy is high, repeat surveillance by cystoscopy yields a high cost for patient health care. The cost of bladder cancer care is estimated at nearly $4 billion per year in the United States of America (USA), which is reported as one of the most costly cancers (*4*). The development of alternative biomarkers is needed to decrease the health cost of recurrence monitoring.

Recent next-generation sequencing technologies have revealed the genetic landscape of human cancers, which supports the idea that tumors acquire cancer-specific somatic mutations in the lineage of mitotic cell divisions (*5*). Through genetic analysis, bladder cancer was revealed to be one of the most frequently mutated cancers, following melanoma, lung cancer, and pancreas cancer (*6*). Theoretically, detecting such gene mutations or their products (mutant proteins) could be expected as 100% cancer-specificity biomarkers.

Evaluating these somatic mutations by circulating tumor DNA in biofluids provides clinical indication for precision therapy through identification of a target molecule (*7*), assessing clonality (*8*), or evaluating the responsibleness of immune checkpoint inhibitors via calculating tumor mutation burden (*9*). Patients with a high tumor burden of the ctDNA showed a high recurrence rate after TUR (*10*). However, the usefulness of detecting recurrence monitoring has not been reported using ctDNA.

Mutant proteins derived from somatic mutant genes are also expected to be promising diagnostic target molecules. However, due to technical factors such as the difficulty in creating antibodies that can specifically recognize a variety of single amino acid substitutions, little progress has been made in their validation. Even with mass spectrometric technology, which can detect a large number of amino acid mutation sequences at once, only a limited number of studies have been reported because the reference sequence database used for identification analysis does not contain information on somatic mutations possessed by individuals. For example, the previous studies reported mutant protein analysis in patient-derived xenograft tumors (*11*) or human colorectal cancer tissues (*12*), but the latter analysis referred to a public comprehensive gene mutation database, which included many mutant sequences that the analyzed patients did not actually have. In this study, we constructed a pipeline to create a database of whole protein sequences, taking into account mutations in each patient, and attempted to identify mutant proteins comprehensively and accurately.

In addition, we focused on extracellular vesicles (EVs) in urine as the resource for detecting mutant proteins. EVs are lipid bilayer membrane vesicles secreted by all cells, which contain nucleic acids and proteins of the original cell and released into body fluids. Since even intracellular proteins that are not secreted proteins are packaged in EVs and can be detected in body fluids, we considered them to be the best samples for detecting bladder cancer-specific mutant proteins in this study. Indeed, multiple functions of mutant proteins in EVs were reported one by one as transforming phenotype of cancer cells (*13*), T cells (*11*), and progressing tumor growth (*14*). However, the comprehensive assessment of mutant proteins encoded by somatic mutations in EVs is absent.

In this work, we established the individual proteogenomic analysis platform and performed comprehensive expression profiling of mutant proteins from bladder cancer tissues, cultured tissue-derived EVs, and urinary EVs. Furthermore, the identified mutant proteins in urinary EVs were evaluated for their usefulness as recurrence monitoring biomarkers using absolute quantitative mass spectrometry techniques and postoperative follow-up urine samples.

## Results

### Construction of the personalized proteogenomic analysis workflow

The experimental overview and concept of the personalized proteogenomic analysis in this study were illustrated in **Figure 1**. To achieve high-confidence identification of non-canonical peptides, such as patient-specific genetic mutation-derived peptides, it is necessary to minimize false positive rate in the database search of mass spectrometric datasets (15). We therefore constructed the individualized whole protein amino acid sequence database for each patient. Following determination of somatic mutations by paired normal/tumor whole exome sequencing, the nucleic acid information was translated to the amino acid one using the Neoantimon package. Here, peptide entries with missense mutations consisted of one mutated amino acid with 25 normal sequences on both sides (**Supplementary Figure 1**). Peptide entries with frameshift mutations consisted of 24 normal amino acids followed by neo-frame sequences until the stop codon (**Supplementary Figure 1**). These mutant peptide entries were added to the human reference sequence database and used in mass spectrometric database searches. In the present study, we comprehensively profiled expression of mutant proteins in bladder cancer tissues, cultured tissue-derived EVs (CTD-EVs), and also urinary EVs from 5 patients who received transurethral resection of bladder cancers (**Supplementary Table 1**).

**Fig. 1.**
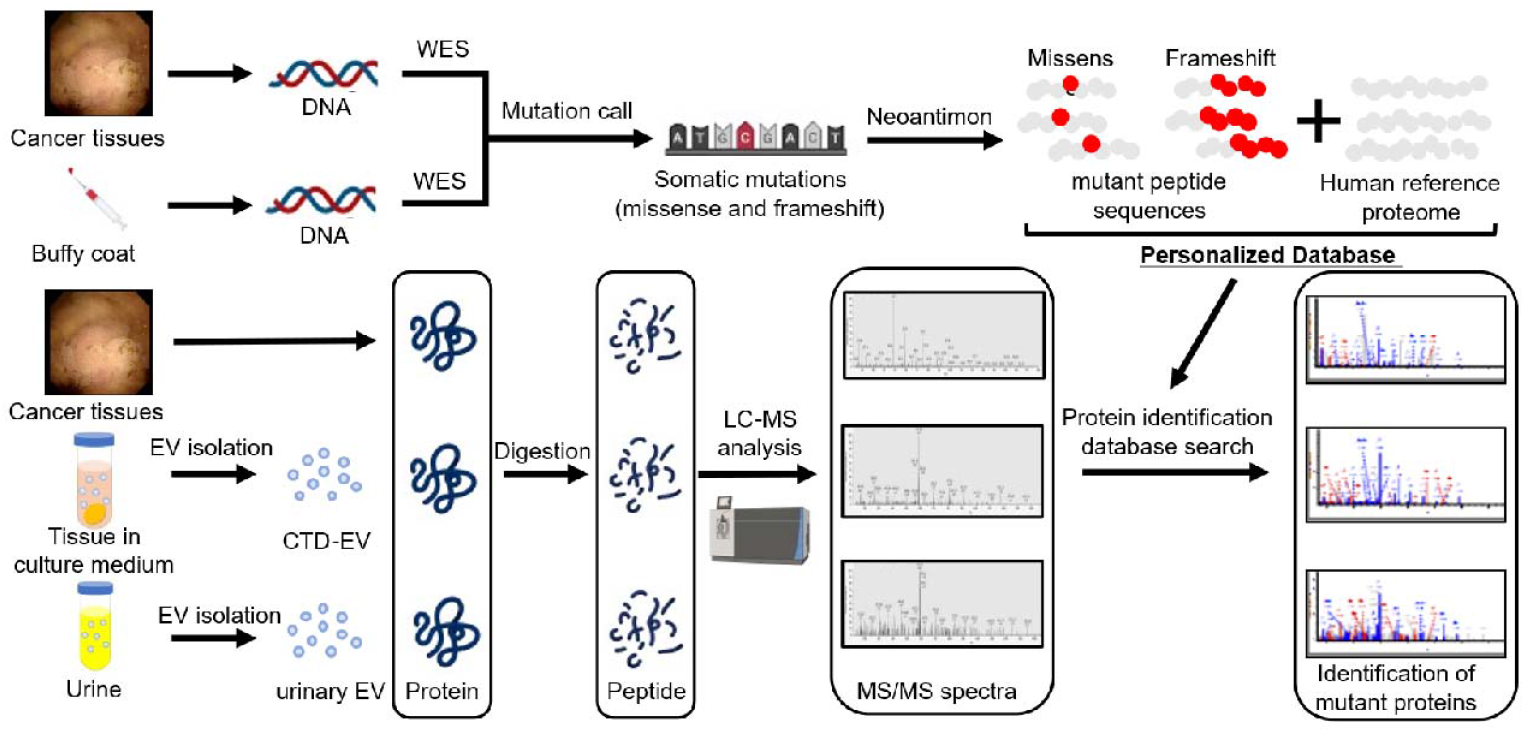
Proteogenomic workflow for identification of mutant proteins. In the upper row, genomic DNA was isolated from cancer tissues or a buffy coat of blood, which was then subjected to whole exome sequencing (WES). Identified missense and frameshift mutations were converted to peptide sequences by Neoantimon. A personalized protein amino acid sequence database (DB) was constructed for each patient. In the lower row, proteins were extracted from frozen cancer tissues, cultured tissue-derived EV (CTD-EV), or urinary EVs. The extracted proteins were digested and analyzed by LC/MS. Here protein identification database search was performed using the personalized protein amino acid sequence database including mutation sequences.

### Identification of somatic mutations using whole exome sequencing

To construct the personalized protein amino acid databases which include mutant sequences, whole exome sequencing (WES) was performed for the first step. From the result of WES for 5 bladder cancer samples, 1,299 single nucleotide variants and 71 frameshift indels were identified in total (**Figure 2A**). The landscape of shared mutated genes among more than one sample is shown in **Figure 2B**. Among these, the somatic mutations of previously reported driver genes such as*TP53, PIK3CA, CDKN1A*, and *ERCC2* were found (*15*). Additionally, frequently mutated genes in whole exome sequencing are detected as *TTN, DNAH1*, and *DNAH10* (*16*).

**Fig. 2.**
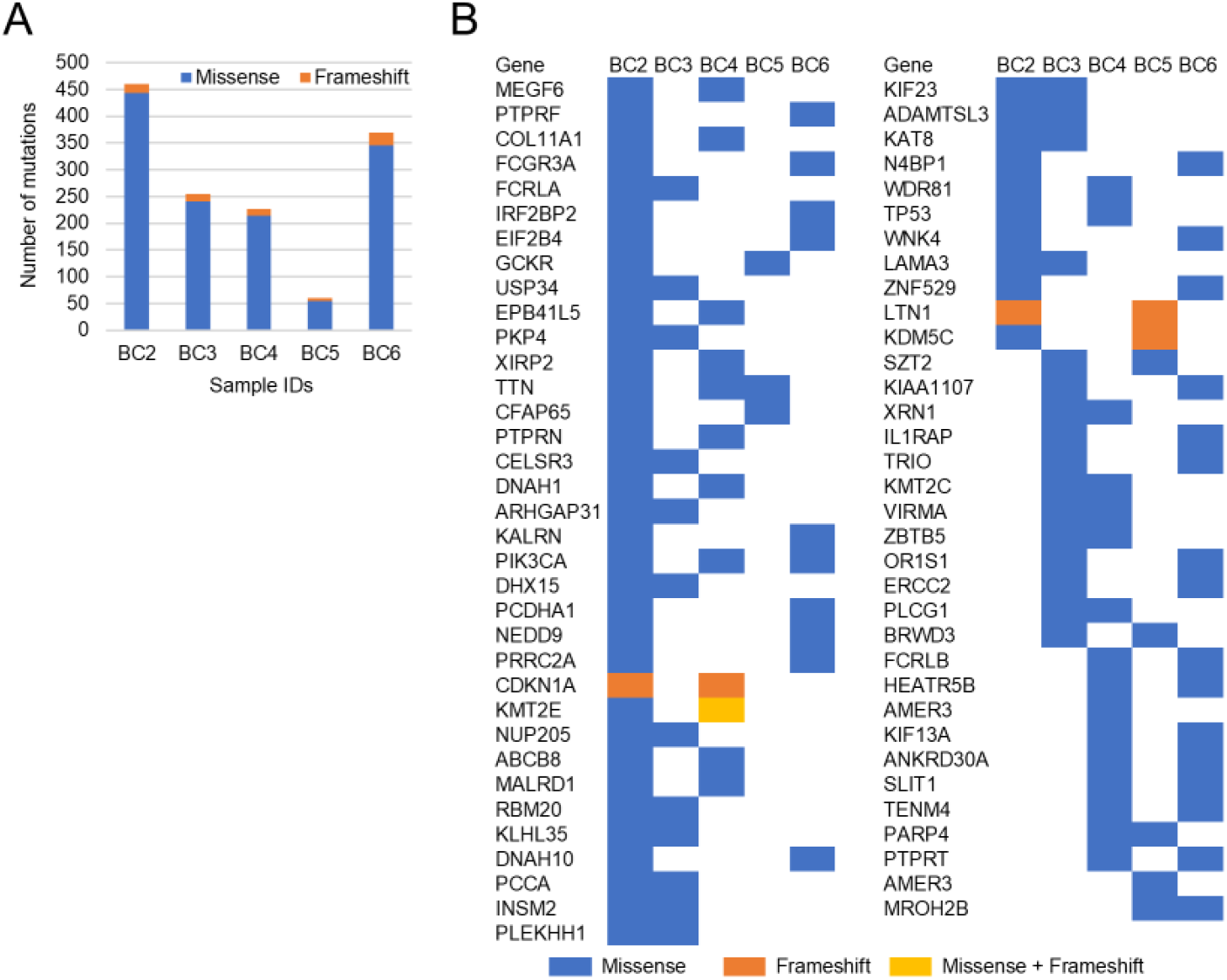
Genetic mutation landscape of bladder cancer tissues. (A) Numbers of missense (blue bars) and frameshift (orange bars) mutations were displayed with the stacked bar chart. (B) List of mutated genes found in more than one tumor sample

### Proteomic landscapes of bladder cancer tissues, CTD-EVs, and urinary EVs

Prior to the proteomic analysis, we assessed quality of EVs isolated from culture medium or urine by use of the Tim4-affinity beads targeting phosphatidylserine on EV surface. The western blotting analysis of crude urine and purified EVs from urine (n = 3) showed remarkably concentrated amount of CD9 and CD63 in the latter samples, which are both known as typical EV marker proteins (tetraspanins) (**Figure 3A**). Similarly, enriched expressions of CD9 and CD63 in CTD-EV samples compared to total cell lysates were confirmed (**Supplementary Figure 2**). In addition, the nanoparticle tracking analysis (NTA) illustrated that median diameter of purified CTD-EVs or urinary EVs was 136 nm or 152 nm, respectively (**Figure 3B**). Based on the previous knowledge, these are also considered as the typical morphological property of EVs (*17*). These datasets (**Figure 3A, 3B, and Supplementary Figure 2**) confirmed that sufficient purity of EVs were isolated from culture media and urine samples. In the present study, bladder cancer tissues, CTD-EVs, and urinary EVs from 5 patients were subjected to quantitative proteome-wide profiling. As the results, 11,207, 9,809, and 5,828 proteins were identified and quantified by LC/MS analyses, respectively (**Supplementary Tables 2-4**), in which 4,686 proteins were commonly detected in the three groups (**Figure 3C**). In comparison with the ExoCarta database which cyclopaedically covers information for the published molecular components of EVs, 57.7% (5,659 proteins) of CTD-EV proteins and 42.8% (2,493 proteins) of urinary EV proteins were newly identified in this study as EV-encapsulated proteins (**Figure 3D**). These data suggested that this study is considered to be one of the highest depth EV proteome analyses derived from clinical specimens compared to previous reports. When evaluating the quantitative correlation between protein contents of bladder cancer tissues and CTD-EVs, weak correlation (R^2^ = 0.241) was found (**Figure 3E**). However, much less correlations were observed in the following pairs, CTD-EVs / urinary EVs (R^2^ = 0.146) and bladder cancer tissue / urinary EVs (R^2^ = 0.097). These values indicated that a group of proteins expressed by bladder cancer tissue could be included in the urinary EVs of bladder cancer patients, but not in a large proportion. Interestingly, however, when the subcellular localization distribution of the three groups was examined, they were almost identical (**Figure 3F**). This means that any protein, including nuclear proteins, can be included in EVs.

**Fig. 3.**
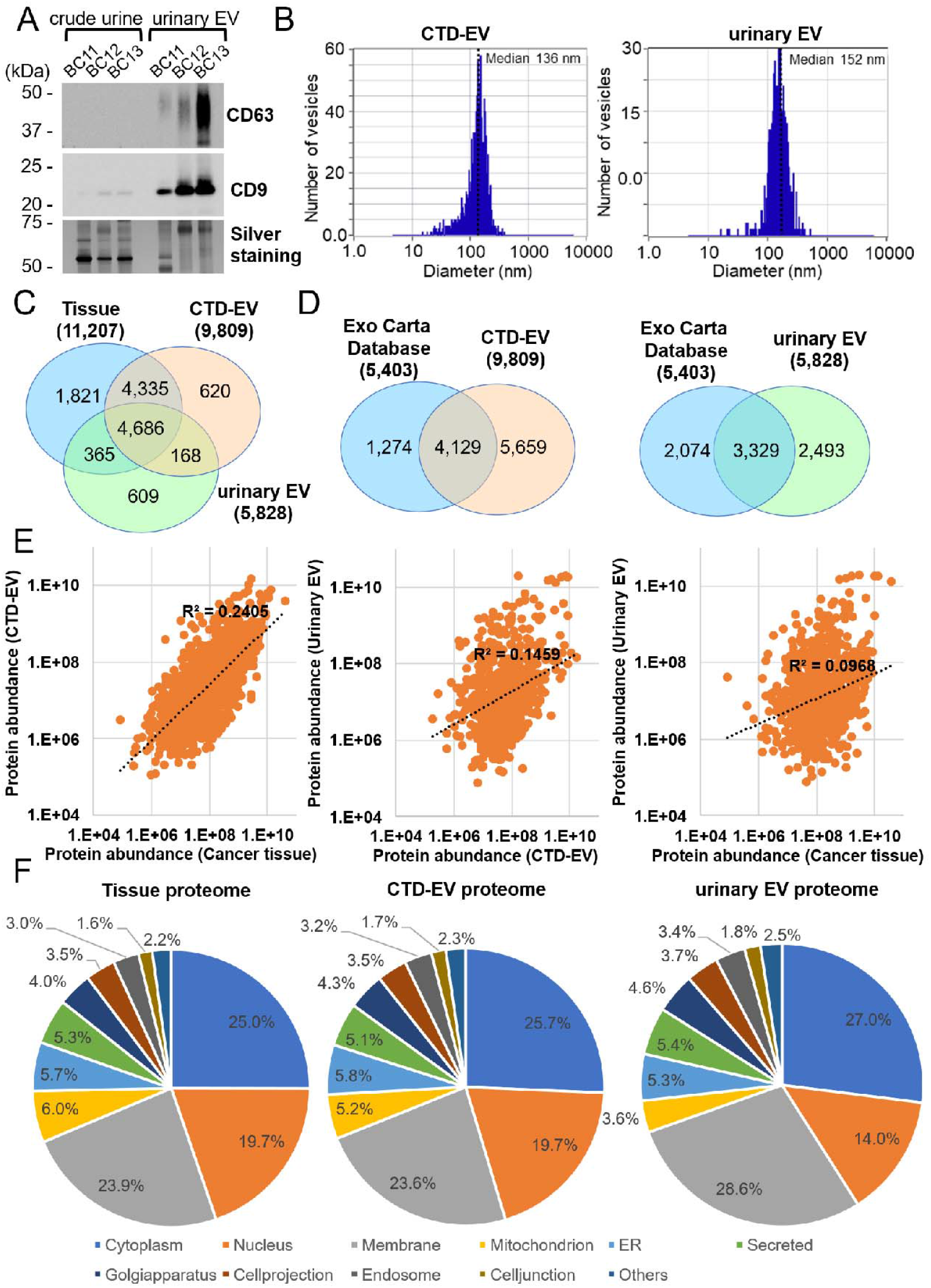
Proteomic landscape of bladder cancer tissues, CTD-EVs, and urinary EVs. (A) Purity of EVs was evaluated by western blotting of EV marker proteins (CD63 and CD9) before (crude urine) or after isolation of EVs (urinary EV) using urine samples from three bladder cancer patients. Total protein was visualized by silver staining. 1 μg of proteins were loaded in each lane. (B) Representative results of nanoparticle tracking analyses which demonstrate distributions of the particle size for CTD-EV or urinary EV. (C) Venn diagram showing numbers of the identified proteins from cancer tissues, CTD-EV, or urinary EV. (D) Comparison of proteins detected in CTD-EVs or urinary EVs with those in the Exo Carta database. (E) Quantitative correlation among tissue proteome, CTD-EV proteome, and urinary EV proteome. (F) Subcellular localizations of the identified proteins from cancer tissues, CTD-EV, and urinary EV.

### Proteogenomic profiling of mutant proteins in bladder cancer tissues, CTD-EVs, and urinary EVs

When broken down into individual patient data, an average of 8,842, 6,418, and 3,374 proteins were detected from cancer tissues, CTD-EVs, and urinary EVs, respectively (**Figure 4A**). Most importantly, these included novel identification of 39, 32, and 4 mutant proteins, respectively, as data for a total of 5 patients (**Figure 4B** and **4C**). Among the 1,299 single nucleotide variants and 71 frameshift indels (**Figure 2A**), protein expressions of 47 single nucleotide variants-containing sequences were confirmed. As a trend, gene mutations with higher variant allele frequencies were more frequently detected as mutant proteins (*p* = 1.3 × 10^−3^, **Figure 4D**). The list of identified mutant peptides from cancer tissues, CTD-EV, or urinary EVs was shown in **Supplementary Table 5, 6**, and **Table 1**, respectively.

**Fig. 4.**
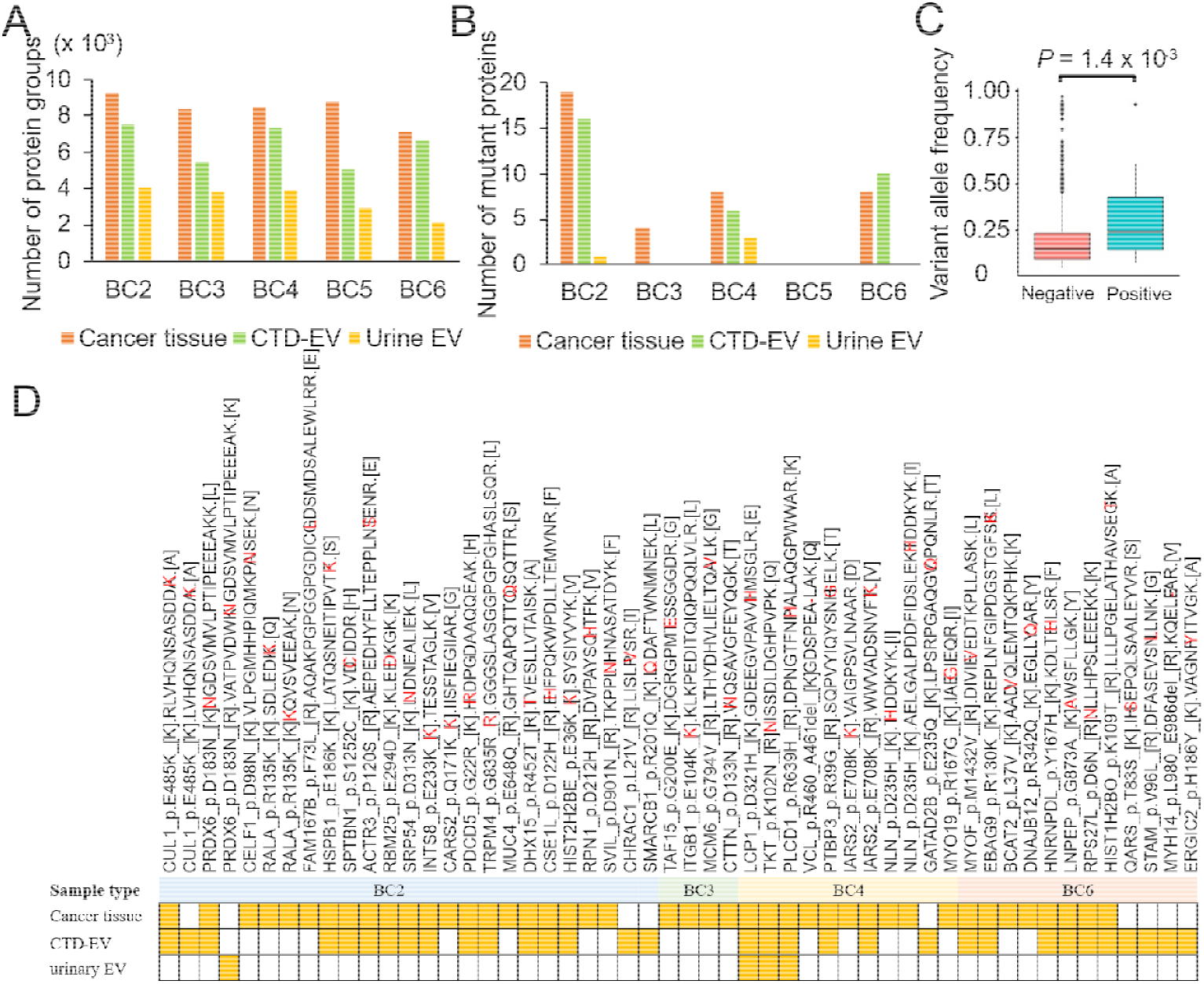
Summary of mutant proteins identified from cancer tissues, CTD-EVs, and urinary EVs. **(A)** The numbers of total proteins identified from cancer tissues, CTD-EV, or urinary EV in each sample were shown. (B) The numbers of mutant proteins identified from cancer tissues, CTD-EV, or urinary EV in each sample were shown. (C) The correlation between positivity of mutant protein detection and variant allele frequency in whole exome sequencing. Negative; genetic variants not detected as proteins, Positive; genetic variants detected as proteins. (D) Comprehensive overview of the mutant proteins detected from cancer tissues, CTD-EV, or urinary EV. The yellow boxes indicate positive detection. The red letters in the amino acid sequences are missense mutations-derived amino acids.

**Table 1.**
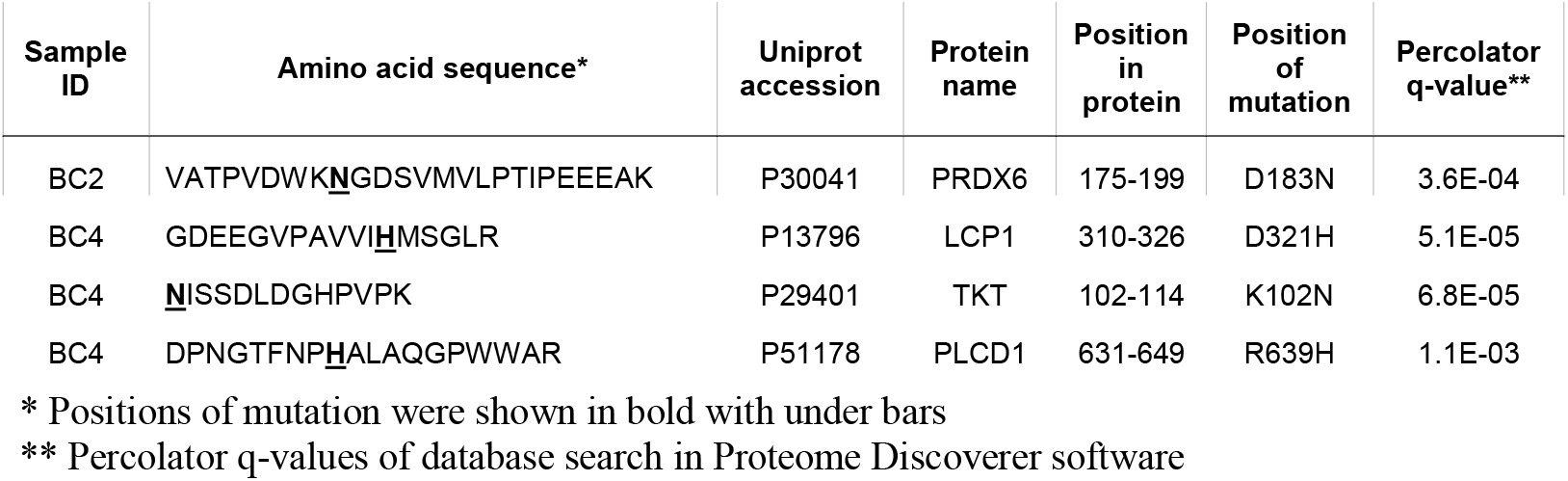
List of identified mutant peptides in urinary EVs.

Among 47 identified mutated amino acid sequences, 11 mutations, CUL1_p.E485K, RBM25_p.E294D, SRP54_p.D313N, SMARCB1_p.R201Q, ITGB1_p.E104K, TKT_p.K102N, PLCD1_p.R639H, IARS2_p.E708K, IARS2_p.E708K, GATAD2B_p.E235Q, and ERGIC2_p.H186Y were previously reported as gene mutations in Catalogue of Somatic Mutations in Cancer (COSMIC) database. Especially, CUL1_p.485K, SMARCB1_p.R201Q, and ITGB1_p.E104K were reported as the gene mutations detected in urinary tract cancer (*17*). However, to our knowledge, this is the first report of detection as mutant proteins for all of the above 47 species.

Regarding the 4 newly identified mutant proteins in urinary EVs, further technical validation was performed using stable isotope-labeled synthetic peptides **(Figure 5** and **Supplementary Figures 3-5**). As shown in **Figure 5A** and the right matrix of **Figure 5B**, the detected *b*- and *y*-series fragment ions covered the entire sequence of endogenous LCP1_310-326_ peptide identified from urinary EVs of the patient BC4, indicating that the amino acid sequence was reliably determined. In addition, the MS./MS spectrum matching between endogenous LCP1_310-326_ peptide (the left panel of **Figure 5B**) and synthetic LCP1_310-326_ peptide (the left panel of **Figure 5C**) demonstrated that the types of fragment ions detected and their intensity patterns matched very well. These data supported that the mass spectrometric identification of LCP1_p.D321H peptide in urinary EVs was convincing. In the similar way, we rigorously confirmed identification of PRDX6_p.D183N, TKT_p.K102N, and PLCD1_p.R639H peptides one by one (**Supplementary Figures 3-5**).

**Fig. 5.**
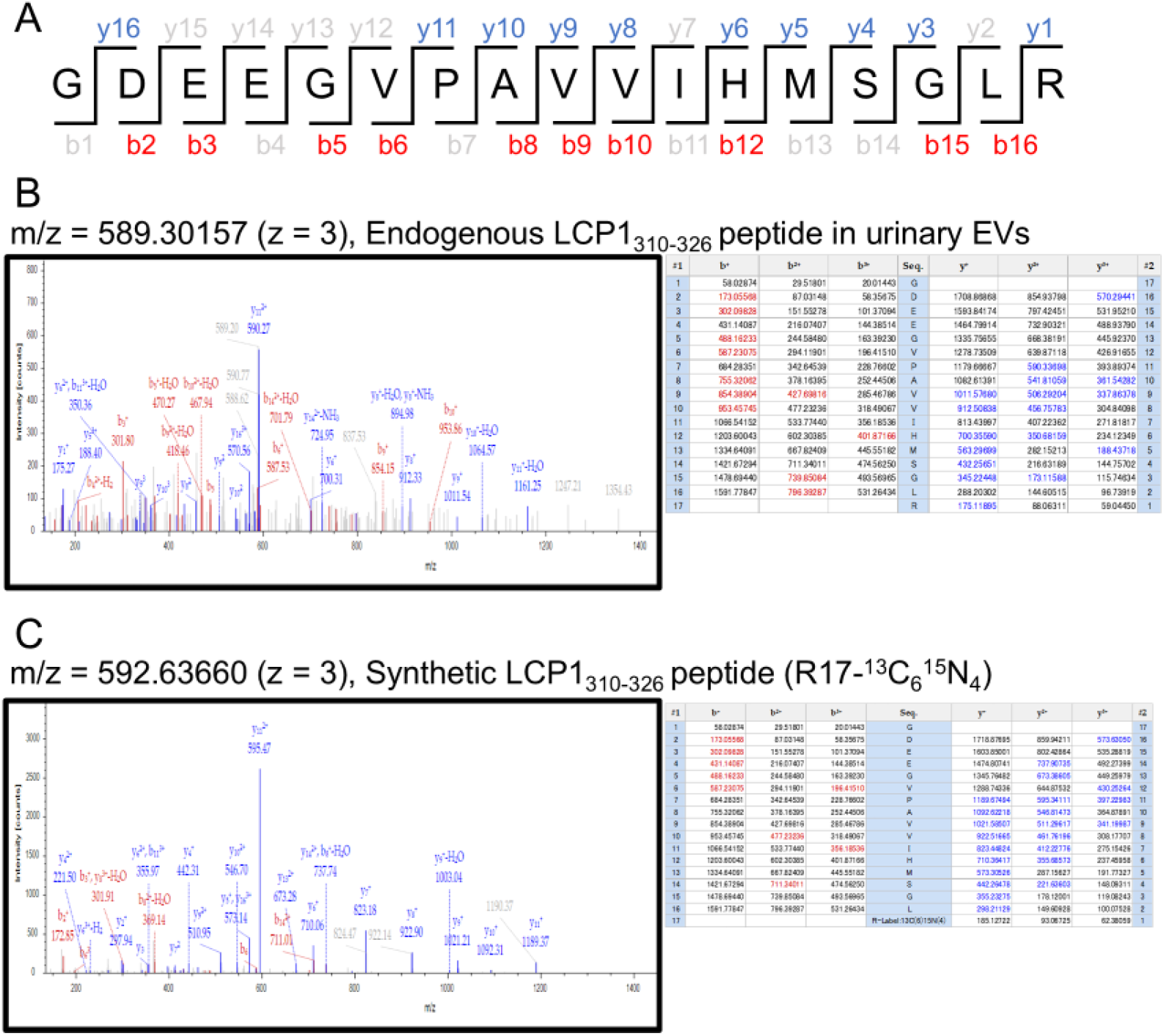
Technical validation of the mass spectrometric identification of LCP1_p.D321H peptide. (A) The amino acid sequence of LCP1_310-326_ peptide is shown, which was detected from the urinary EV sample of the patient BC4. The b- and y-series fragment ions identified in the Sequest database search were displayed in red and blue letters, respectively. The centroid MS/MS spectrum of endogenous LCP1_310-326_ peptide detected from urinary EVs (B) or synthetic LCP1_310-326_ peptide (C) was shown in the left panel. The right matrix demonstrates the sequence coverage in the Sequest database search. The b- and y-series fragment ions identified in the Sequest database search were displayed in red and blue letters, respectively.

### A diagnostic potential of mutant proteins in urinary EVs for recurrence monitoring of bladder cancer

To assess the diagnostic potential of EV mutant proteins as a non-invasive urine test, the absolute quantification measurement methods were established for the three mutant proteins detected in urinary EVs of the patient BC4 (LCP1_p.D321H, TKT_p.K102N, and PLCD1_p.R639H). Here we prepared the following synthesized peptides as absolute quantification standards, LCP1_p.D321H_310-326_, TKT_p.K102N_102-114_, and PLCD1_p.R639H_631-649_ in which C-terminal Arg or Lys was replaced to Arg-^13^C_6_ ^15^N_4_ or Lys-^13^C_5_ ^15^N_3_, respectively. We optimized collision energy in peptide fragmentation and compensation voltage (CV) of field asymmetric ion mobility spectrometry (FAIMS) in parallel reaction monitoring (PRM) mode to maximize the detection sensitivity for each of the three peptides (**Supplementary Figure 6**). Using the optimized PRM methods, presurgical and postsurgical (3 or 6 months after surgery) urine samples collected from the patient BC4 (**Figure 6A**) were analyzed. The PRM chromatograms in **Figure 6B** demonstrated that the signal of LCP1_p.D321H_310-326_ was clearly diminished in the postsurgical urinary EVs even 6 months after surgery. Interestingly, the absolute concentrations of all three mutant proteins, LCP1_p.D321H, TKT_p.K102N, and PLCD1_p.R639H, were significantly reduced in the postoperative urinary EV samples (**Figure 6C**). More importantly, the results of this urinalysis were well consistent with the diagnosis of no recurrence on cystoscopy at 6 months postoperatively.

**Fig. 6.**
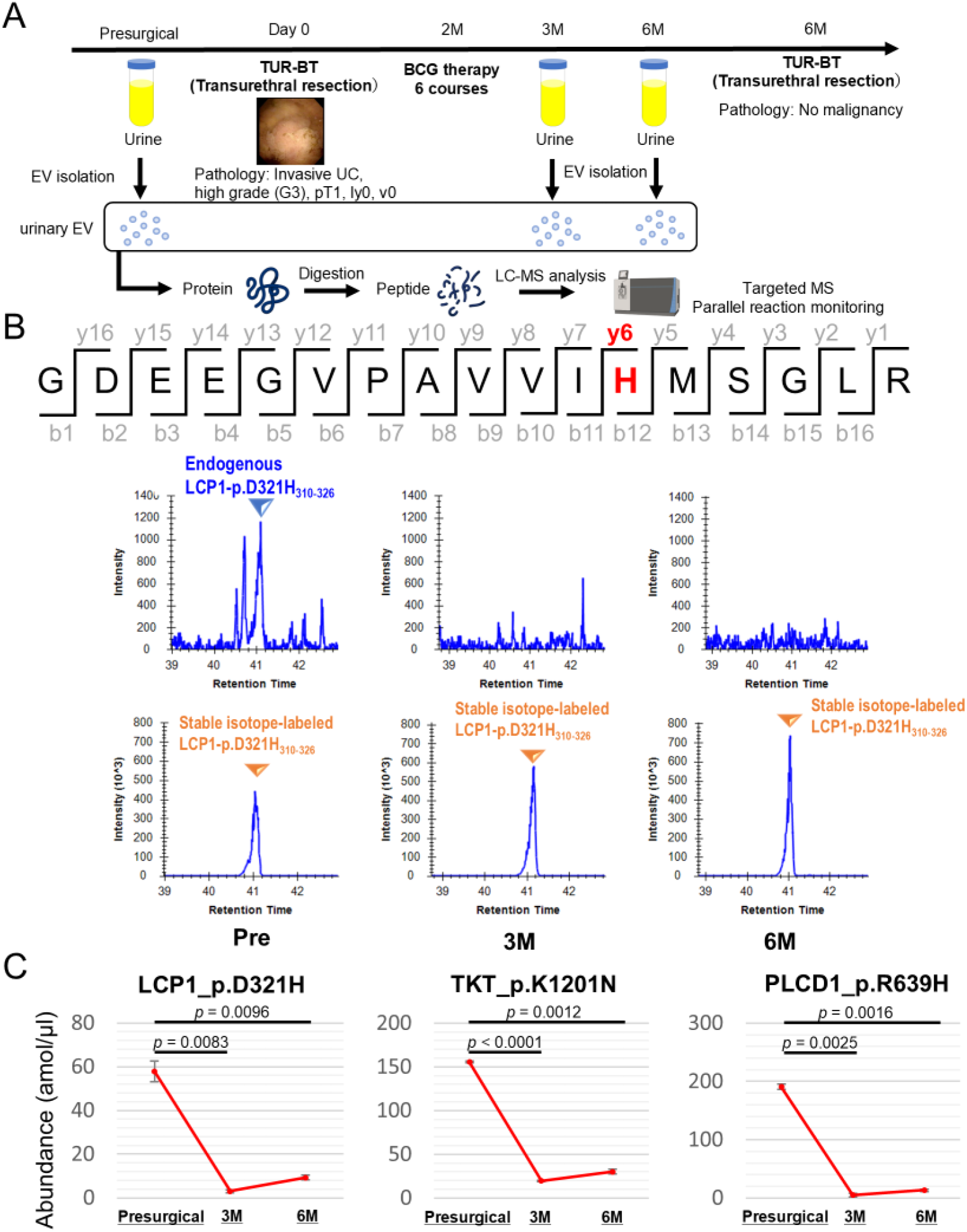
Recurrence monitoring of bladder cancer using mutant proteins in urinary EV. (A) Timeline of sample collection and flow chart of experimental procedures. The patient received TURBT and had no recurrence at 6 months after surgery. EV was isolated from urine samples collected at presurgical point and 3 or 6 months after surgery. (B) The upper row shows amino acid sequence of LCP1-p.D321H_310-326_, in which the missense mutation-derived amino acid His is displayed in red. The 6 panels in the lower rows are PRM chromatograms of the y6 product ion derived from LCP1-p.D321H_310-326_ peptide detected in urinary EV at presurgical point (left) and 3 (center) or 6 months after surgery (right). 10 fmol of the stable isotope-labeled standard peptide (bottom row) was spiked into samples and simultaneously measured with the intact peptide (middle row). (C) The abundance of mutant peptides LCP1_p.D321H, TKT_p.K120N, and PLCD1_p.R639H in urinary EVs of patient BC4 were shown at the three time points above. The *p*-values were determined by the two-sided parametric student t-test. The data showed the mean ± SD (n = 3).

## Discussion

Although a vast number of gene mutations have been analyzed in many types of cancer, including bladder cancer, comprehensive knowledge of which of them are expressed as mutant proteins, and in what quantities, has rarely been reported. One example of the comprehensive mutant protein analysis was the study combining RNA sequencing with mass spectrometric analysis using 86 colorectal cancer tissues (*12*). Here, 108 mutant proteins were detected in 25 cases. However, in the hyper-mutated group (more than 12 mutations per megabase), mutant proteins were detectable in 81% of cases, while in the non-hyper-mutated group, mutant proteins were detectable in only 12% of cases. In contrast, in the present study, mutant proteins were detected in four cases from five bladder cancer tissue samples. The reason for the higher detection rate of mutant proteins can be attributed to the construction and use of an individualized protein sequence database. In contrast to the previous report, in which protein identification database searches were performed against the protein sequence database pooled from comprehensive RNA sequencing data for 86 cases, in our study, a protein sequence database was created and searched for each individual mutation. We believe that this minimized the false positive rate and supported identification of rare mutant proteins. In addition, the depth of proteome analysis itself has been improved. In fact, we succeeded in detecting 11,207 proteins from 5 tissue samples, whereas 7,526 proteins were detected from 86 tissue samples in a previous report.

On the other hand, significant progress has been made in the analytical depth of urinary EV proteins compared to previous reports. As a recent example, there is a report that 1,960 proteins were detected in the proteome analysis of urinary EVs (*23*), while 5,828 urinary EV proteins were successfully identified in this study. This can be attributed to two factors. First, an extremely high degree of EV purification was achieved. In this study, Tim4 protein-coated magnetic beads, which specifically capture phosphatidylserine expressed on EV membranes (*22*), were used, and in our experience, these beads can produce overwhelmingly higher purity EVs than any other EV purification methods, including ultracentrifugal methods, size exclusion chromatography, and so on. In fact, as data, considerably more EV marker proteins were detected (CD9, CD63, CD81, CD82, CD151, TSG101, ALIX, Tetraspanin-9, 13, 17, XX, XX, and XX) compared to previously reported proteome analyses of EVs obtained by other purification methods (**Supplementary Tables 3 and 4**). Second, FAIMS was optimized to maximize EV protein identification. In addition to liquid-phase separation by HPLC, real-time gas-phase separation by FAIMS was added, which allowed even greater depth of protein detection than conventional LC/MS analysis.

There are as yet no comprehensive reports of mutant proteins present in EVs. Only few studies investigated for specific major driver gene products in cancer cell-derived EVs. For example, an antibody-based single EV analysis technique was reported (*30*). In that study, EVs were isolated from serum using size exclusion chromatography and stained with fluorescently labeled antibodies against mutant KRAS and TP53, followed by spread on glass slides for image detection (*30*). By use of this method, mutant KRAS and TP53-positive EVs were detected from sera of patients with early stage pancreatic cancer. However, such a technique cannot detect a panel of many species like ctDNA because it is extremely difficult to create good antibodies specific for single amino acid substitutions. In the recurrence monitoring experiment (**Figure 6**), we demonstrated that three mutant proteins in urinary EVs were simultaneously and absolutely quantified in a single PRM analysis. Thus, the advantage of the mass spectrometry-based detection method is the ability to simultaneously measure multiple items without change in cost, regardless of antibody availability. In this study, 32 or 4 mutant proteins were detected in CTD-EVs or urinary EVs, respectively, however, when the number of cases measured would be expanded in the future, more mutant proteins in EVs could be found, leading to establishment of a mutant protein panel test covering more patients in a single analysis.

In actual bladder cancer treatment, cystoscopy is required every few months over a long period of time for recurrence monitoring, but its high invasiveness is problematic. Although urine cytology is a typical urine-based non-invasive test with high specificity, sensitivity is insufficient. For molecular biomarkers in urine, nuclear matrix protein 22 (NMP22), bladder tumor antigen (BTA), and uroVysion have been approved as clinical tests, but their specificity is lower than that of urine cytology (*19-21*), and they are rarely used as a priority test. In this study, we showed that the quantitative values of mutant proteins in urinary EV can reflect the cystoscopy findings, which are non-invasive and have, in principle, 100% cancer specificity. Therefore, mutant proteins in urinary EV could be expected as a new urine liquid biopsy test that can avoid frequent cystoscopy.

Considering the biological significance of mutant proteins in EVs, the fact is that there are still many aspects that are not well understood. The 47 mutant proteins we identified in bladder cancer tissues included CUL1_p.E485K, SMARCB1_p.R201Q, TKT_p.K102N, PLCD1_p.R639H, and IARS2_p.E708K, which were reported to be frequently mutated in bladder cancer. Among these, an interesting fact is known about the relationship between CUL1 mutations and cancer. CUL1, together with Skp1 and F-box proteins, forms the SCF ubiquitin ligase (E3) complex, which mainly degrades proteins involved in cell cycle progression (p27_Kip1, Cyclin E, etc.). Remarkably, more than 50% of CUL1 mutations reported in various cancer types involve amino acid substitutions from Glu to Lys, such as E420K, E485K, E493K, and E733K (*18*). These substitutions are thought to have the effect of partially reversing the charge on the C-terminal region of CUL1, resulting in enhancing binding of the scaffold protein CUL1 to other complex-forming proteins, leading to excessive degradation of cell cycle-related proteins and hyperproliferation of cells. The function of CUL1_p.E485K in EVs has not yet been determined, but it is possible that this mutant protein propagates to surrounding cells via EVs and contributes to cell cycle disruption in recipient cells. In fact, some mutant proteins such as EGFR vIII and KRAS_G13D are known to be encapsulated in EVs and transferred to function in immune cells and other cells in the cancer microenvironment (*13, 14*). Regarding LCP1_p.D321H, PRDX6_p.D183N, TKT_p.K102N, and PLCD1_p.R639H detected in urinary EV, it is known that LCP1 is responsible for cell motility, PRDX6 is a multifunctional enzyme involved in cell oxidative stress response and lipid metabolism, TKT is a key enzyme in non-oxidative pentose phosphate pathway, and PLCD1 is a member of the phospholipase C family that plays a key role in intracellular signal transduction, however, the functional changes caused by the above amino acid substitutions have not been reported.

In conclusion, we established a novel liquid biopsy technique to detect mutant proteins in urinary EVs that could be useful for monitoring bladder cancer recurrence. These, like ctDNA, are thought to have extremely high cancer specificity, and their further development (ctDNA and ctProtein) will complementarily improve the accuracy of bladder cancer diagnosis. In addition, it will be necessary to expand the mutant protein panel more, such as the driver mutant proteins, to cover a wider population.

## Materials and Methods

### Sample collection

Presurgical urine, cancer tissues, blood, and postsurgical urine were collected from 5 patients who received transurethral resection of bladder cancer at the University of Tokyo Hospital. Fresh tumors were frozen in liquid nitrogen immediately after surgical resection. All tumor tissues were diagnosed as urothelial cancer by the pathologist. This study was performed under the Declaration of Helsinki and approved by the ‘Ethics Committee of the Cancer Institute Hospital of Japanese Foundation for Cancer Research’ (approval number #2022-GB-055) and ‘Ethics Committee of the University of Tokyo Hospital’ (approval number 2022353Ge), and all the patients in the present cohort provided written informed consent before the study entry.

### Whole exome sequencing and constructing individual protein amino acid sequence database

Genomic DNA was extracted from cancer tissues and buffy coats of the 5 bladder cancer patients. Following the manufacturer’s protocol, DNA was isolated from 2 to 10 mg of frozen tissues or buffy coat from 8.5 ml of blood using the QIAamp DNA Mini Kit (QIAGEN). Blood was collected using a cell-free DNA collection tube (Roche). DNA library preparation with SureSelect Human All Exons V6 kit (Agilent Technology) and sequencing with the Illumina Novaseq 6000 were performed. Sequencing reads were aligned to the human genome hg 19, and somatic nonsynonymous and frameshift mutations were identified with the Genomon 2 pipeline (https://genomon.readthedocs.io/ja/latest/). Somatic mutations were filtered by the following criteria: (i) number of variant reads in tumor > 3, (ii) number of variant reads in normal < 3, (iii) variant allele frequencies in tumor > 0.05, and (iv) excluding simple repeat (*33*). Nonsynonymous and frameshift mutations were extracted by the Neoantimon R package software to construct a customized database of the mutant proteins (*34*). This software generates mutant peptides by constructing the corresponding RNA sequences using the Reference Sequence Database (*34*). Finally, a customized personalized database was constructed for each patient by combining the mutant peptides sequence from the Neoantimon software and the reference sequence from Swiss-Prot human proteome database (20,386 entries).

### Protein extraction from tissues

The frozen cancer tissues were lysed with 100 µL of lysis buffer [20 mM HEPES (pH 7.8), 150 mM NaCl, 1% NP-40, 0.1% SDS, 10% glycerol, and cOmplete Protease Inhibitor Cocktail (Sigma-Aldrich)] on ice. The tissues were homogenized using BioMasher II (Funakoshi). After centrifugation at 15,000 × *g* for 15 min, the supernatant was collected. Protein concentration was measured using the Micro BCA Protein Assay Kit (Thermo Fisher Scientific).

### Isolation of cultured tissue-derived EV and urinary EV

Cultured tissue-derived EV (CTD-EV) was isolated from culture medium of cancer tissues. First, the 2-mm cubes of the frozen cancer tissues were thawed on ice and rinsed with PBS and incubated in 1.5 mL of RPMI-1640 with gentle rotation at 37 °C for 3 hours. Here, the culture medium was supplemented with 10% EV-depleted fetal bovine serum (FBS) (Funakoshi) and 1% penicillin-streptomycin. After centrifugation at 3,000 × *g* for 5 min, the supernatant was stored at -80 °C until use. Urine sample was centrifuged at 2,000 × *g* for 10 min and the supernatant was stored at -80 °C until use. Urine was concentrated to 1 ml using AmiconUltra 100K molecular weight cutoff concentrators (Merck) before EV isolation. EVs were isolated from culture media of tissues or 5 ml of urine using the MagCapture Exosome Isolation Kit PS Ver.2 (FUJIFILM Wako Pure Chemical Corporation) according to the manufacturer’s instruction. Only for the proteome analysis, EVs were eluted with 30 *μ* l of the Laemmli’s SDS sample buffer.

### Western blotting and silver staining analyses

The isolated proteins (10 µg) in CTD-EV and urinary EV from bladder cancer patients were separated on 10% SDS polyacrylamide gels (Thermo Fisher Scientific) and transferred onto membranes. Membranes were blocked with 4% block Ace (KAC), followed by incubation with the anti-CD9 (SHI-EXO-MO1, Cosmo Bio) or anti-CD63 (SHI-EXO-MO1, Cosmo Bio) antibodies. After washing, the membranes were incubated with HRP-conjugated anti-mouse IgG antibody (NA9310V, GE Healthcare). Finally, protein signals were detected with Western Lightning ECL Pro (Nacalai tesque) reagent and ChemiDoc imaging system (Bio-Rad). Silver staining of the gels was performed using the SilverQuest Silver Staining Kit (Invitrogen) according to the manufacturer’s instruction.

### Nanoparticle Tracking Analysis

CTD-EV and urinary EV isolated with the MagCapture Exosome Isolation Kit PS Ver.2 were diluted 1: 50 in PBS. The size distribution of EVs was analyzed using Zetaview (Particle Metrix) with analysis parameters: Maximum area: 1000, Minimum area 5, Minimum brightness: 30, and Laser wavelength: 488 nm.

### Sample preparation for mass spectrometric analysis

Proteins were digested using the S-Trap micro columns (ProtiFi) according to the manufacturer’s instruction. Here, Trypsin/LysC Mix (Promega) in 20 µl of 50 mM TEAB was used for digestion reaction at 47 °C for 2 hours. The eluted peptides were dried by Speed-Vac sample concentrator and stored at -30 °C before LC-MS analysis.

### LC/MS analysis

The digested peptides were resuspended in 2% acetonitrile with 0.1% formic acid and analyzed using Orbitrap Lumos Fusion Tribrid mass spectrometer equipped with High-Field Asymmetric-waveform Ion Mobility Spectrometry (FAIMS) Pro interface (Thermo Scientific) combined with Vanquish Neo UHPLC system (Thermo Scientific). The sample was trapped by a precolumn (Pepmap Neo C18 5 µm, 300 µm × 5 mm, Thermo Scientific). The trapped sample was separated by analytic column (Aurora UPHLC Column, C18, 0.075 × 250 mm, 1.6 µm FSC, IonOpticks). Mobile phase A consisted of 0.1% formic acid and mobile phase B consisted of 0.1% formic acid in acetonitrile. The Vanquish Neo system was used at a flow rate of 200 nl/min using a linear gradient starting from 2% to 30% B by 115 min, followed by increase to 95%B by 117 min and held for 3 min before returning to 2% B. The compensation voltages (CVs) were set at [-40, -60, and -80V] for set-1, [-50, -70, and -90] for set-2, and [-45, - 55, and -65] for set-3. A full MS scan was collected from m/z = 350-1,500 at a resolution of 120K. The maximum injection time for the full MS scan was 50 ms with auto gain control (AGC) of 4e5. Charge states 2 to 5 were selected for fragmentation by higher energy collisional dissociation (HCD) at the 30% normalized collision energy. The maximum ion injection time for the MS2 scan was 22 ms with an AGC of 3e4 by ion trap.

### Protein identification analysis

Global peptide identification including mutant peptides was performed using customized personalized database using Proteome Discoverer version 3.0 (Thermo Scientific). The obtained raw files were processed by spectrum selector, followed by peptide identification by Mascot and Sequest HT. Sequest search was combined with precursor detector, spectrum properties filter, and INFERYS rescoring. One or two missed cleavages were allowed for searching tryptic peptides. Carbamidomethylation of cysteine was set as static modification and oxidation of methionine was set as dynamic modification. The precursor and fragment mass tolerance were set to 10 ppm and 0.6 Da, respectively. False discovery rate (FDR) was set to 0.01 (Strict) to identify highly confident peptides using percolator.

### Validation of identified mutant peptides

Stable isotope-labeled peptides were synthesized and used for further confirmation of identified mutant peptides in urinary EV (**Table 1**). The C-terminal lysine (Lys) or arginine (Arg) were replaced with Lys-^13^C_3_ ^15^N_5_ or Arg-^13^C_4_^15^N_6_, respectively. The synthetic peptides were spiked to the peptide samples at the final concentration of 100 fmol/injection.

### Absolute quantification by parallel reaction monitoring

For absolute quantification of the mutant proteins in urinary EVs, parallel reaction monitoring (PRM) was performed using Orbitrap Fusion Lumos Tribrid mass spectrometer equipped with FAIMS Pro interface combined with Vanquish Neo UHPLC system. The tMS2 node in the Xcalibur software (Thermo Scientific) was used for PRM acquisition. The PRM parameters, including m/z and CV, for the three mutant peptides (LCP1_p.D321H, TKT_p.K102N, and PLCD1_p.R639H) were shown in **Supplementary Table 7**. Peptide abundance was calculated using the area under the curve of endogenous and isotope-labelled peptide peaks. Detection of endogenous and isotope-labeled peptide peaks was performed automatically by Skyline software (*35*).

## Statistical analysis

Statistical analysis was performed using the statistical software R 4.4.0 version. Abundances of mutant proteins in **Figure 6** were displayed as the mean with standard error. The *p*-values were calculated by paired Student’s test.

## Data availability

The whole exome sequencing and LC/MS raw data were deposited in the Japan Proteome Standard Repository/Database (jPOST), JPST003684.

## Author contributions

Conceptualization: Y.H, K.U.; Experiments: Y.H, K.S., H.S.; Clinical samples: Y.H., K.S., Y.Y., T.D., K.H.; writing – original draft; Y.H., K.S; writing – review and editing: Y.Y., H.K., K.U.

## Data and materials availability

All data needed to evaluate the conclusions in the paper are present in the paper and the Supplementary Materials.

## Supplementary Materials

This PDF file includes: Figs. S1 to S6

Legends for tables S1 to S7

Other Supplementary Material for this manuscript includes the following:

Tables S1 to S7

