## Supplementary Materials for "Development of a liquid biopsy for bladder cancer using a mutant protein panel in urinary extracellular vesicles"

Example: Yuji Hakozaki *et al.*

**This PDF file includes:**

Figs. S1 to S6

Legends for tables S1 to S7

**Other Supplementary Material for this manuscript include the following:**

Table S1 to S7

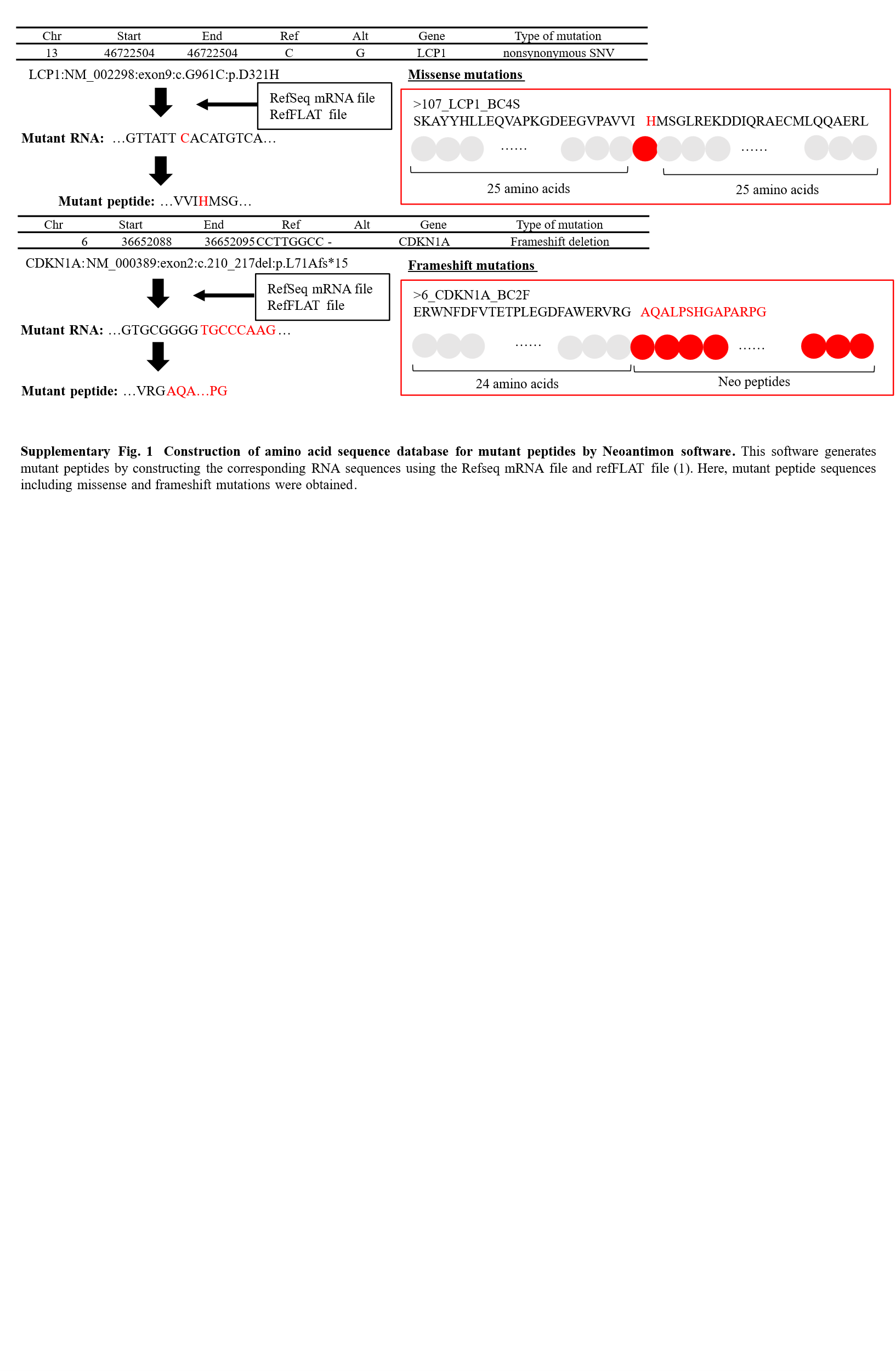

Fig. S1. Construction of amino acid sequence database for mutant peptides by Neoantimon software. This software generates mutant peptides by constructing the corresponding RNA sequences using the Refseq mRNA file and refFLAT file. Here, mutant peptide sequences including missense and frameshift mutations were obtained.

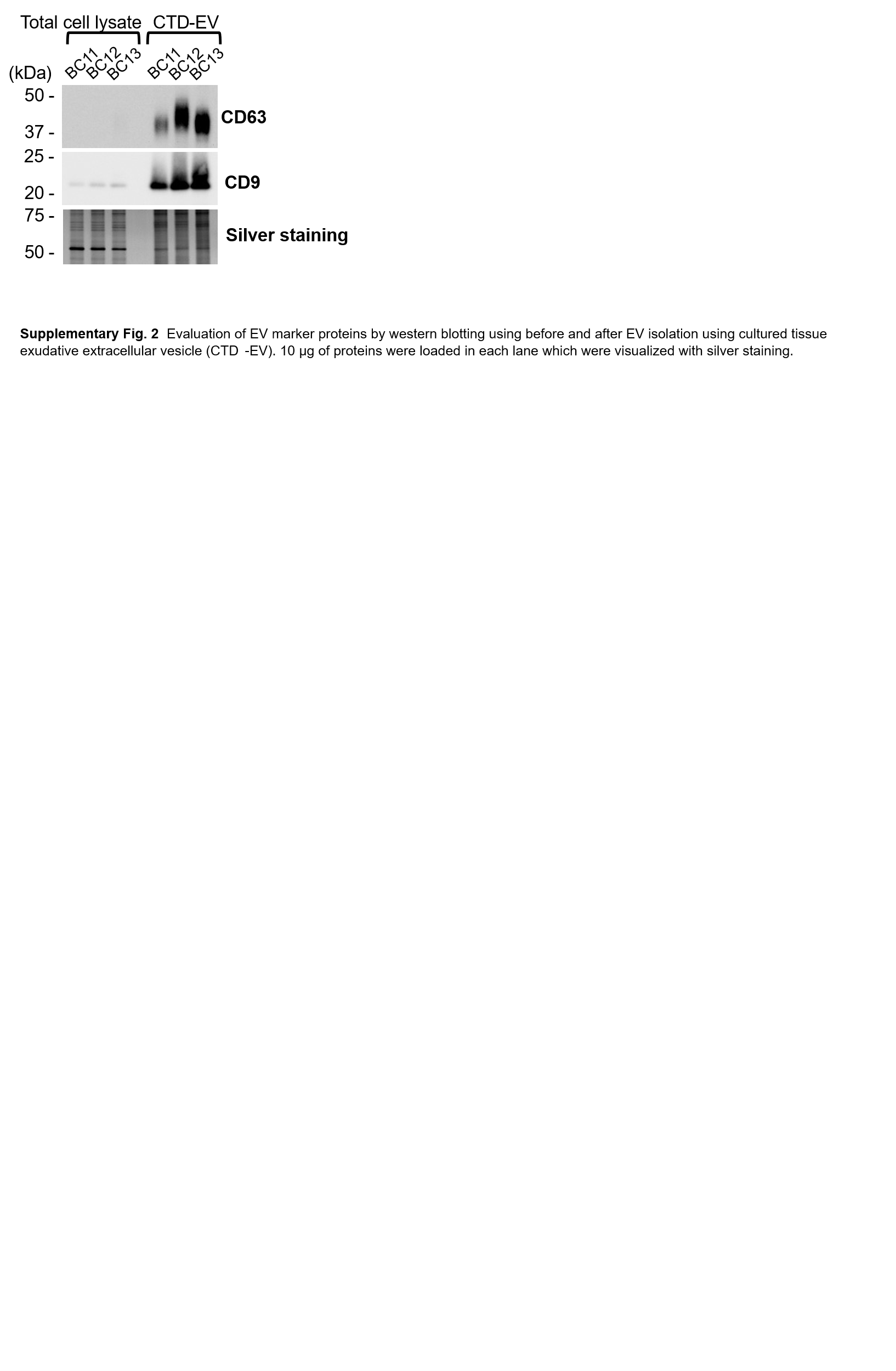

Fig. S2. Evaluation of EV marker proteins by western blotting using before and after EV isolation using cultured tissue exudative extracellular vesicle (CTD-EV). 10 μg of proteins were loaded in each lane which were visualized with silver staining.

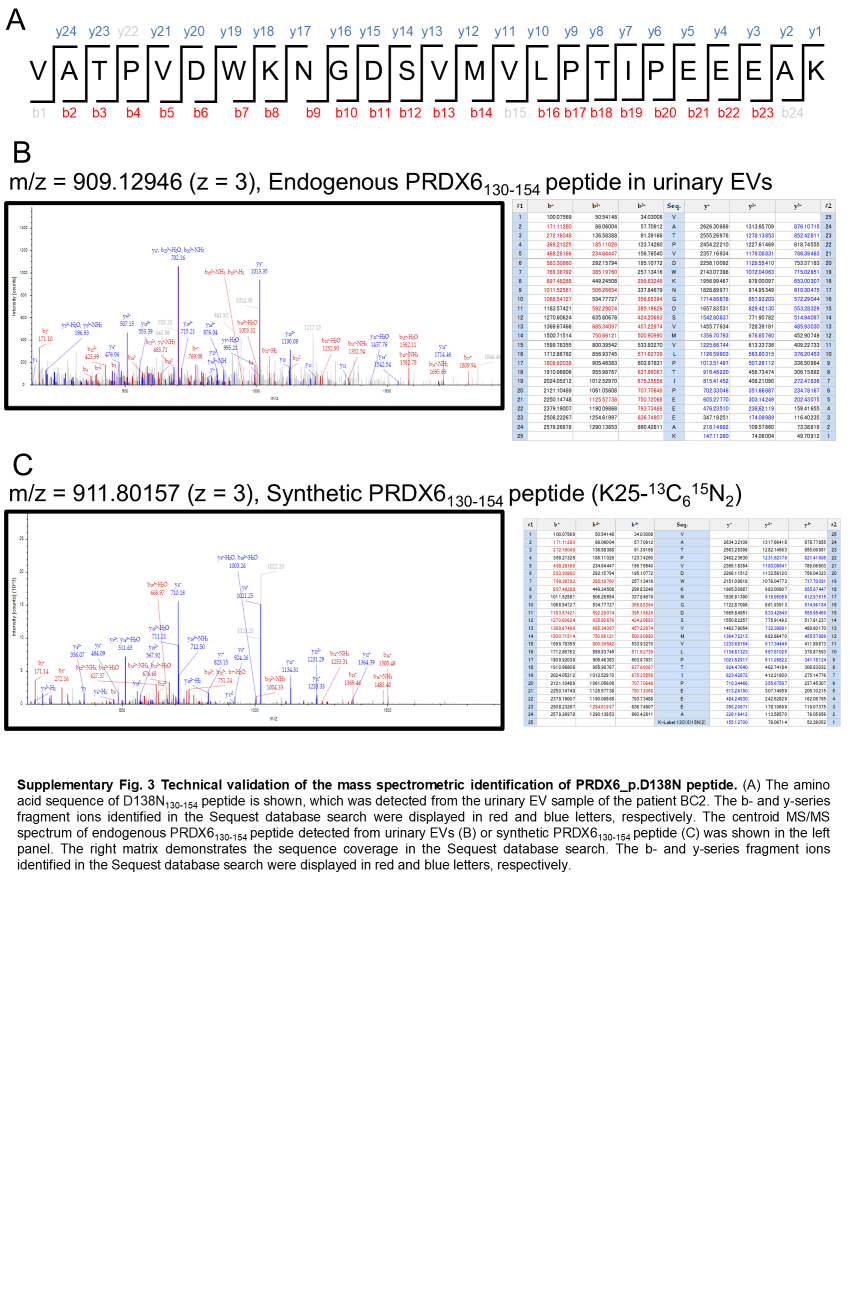
Fig. S3. Technical validation of the mass spectrometric identification of PRDX6_p.D138N peptide. (A) The amino acid sequence of D138N_130-154_ peptide is shown, which was detected from the urinary EV sample of the patient BC2. The b- and y-series fragment ions identified in the Sequest database search were displayed in red and blue letters, respectively. The centroid MS/MS spectrum of endogenous PRDX6_130-154_ peptide detected from urinary EVs (B) or synthetic PRDX6_130-154_ peptide (C) was shown in the left panel. The right matrix demonstrates the sequence coverage in the Sequest database search. The b- and y-series fragment ions identified in the Sequest database search were displayed in red and blue letters, respectively.

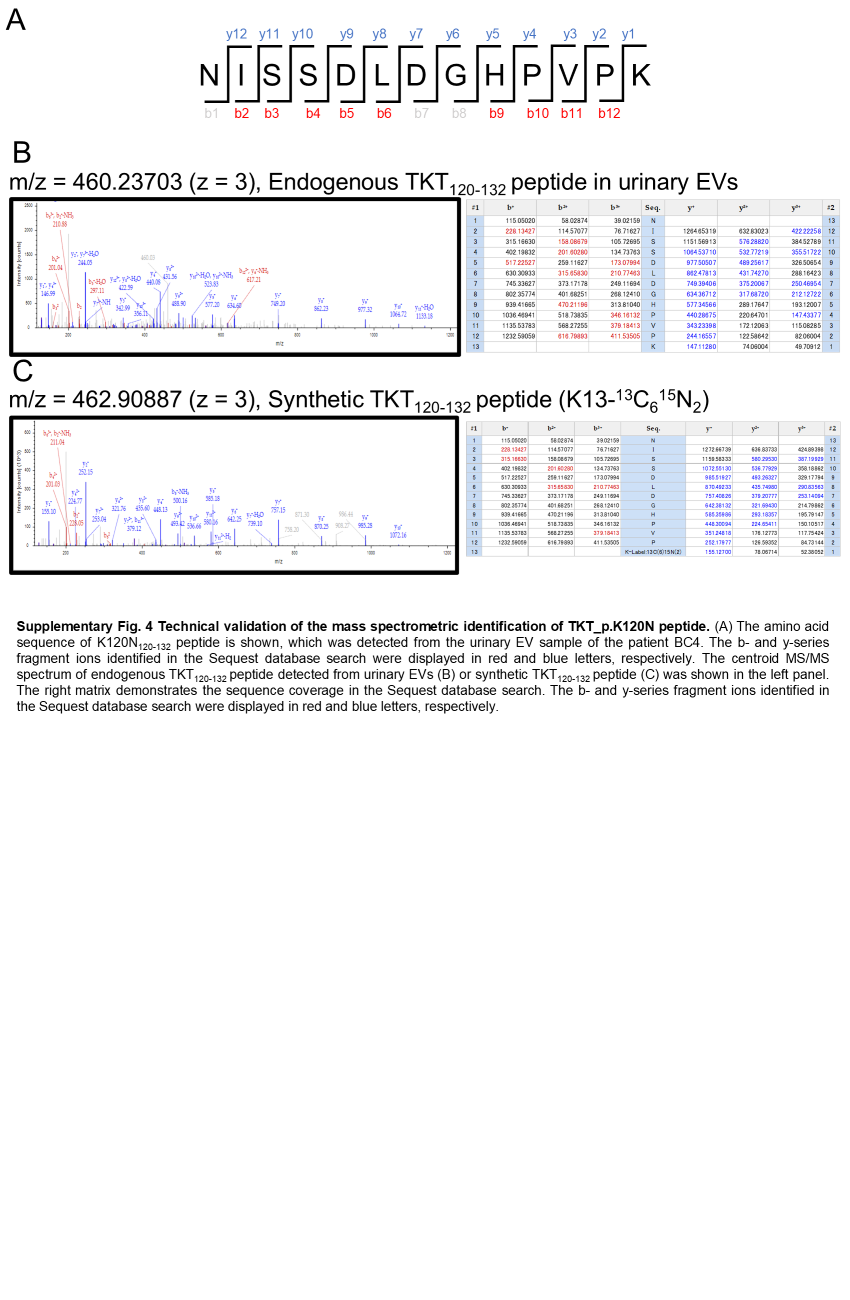
Fig. S4. Technical validation of the mass spectrometric identification of TKT_p.K120N peptide. (A) The amino acid sequence of K120N_120-132_ peptide is shown, which was detected from the urinary EV sample of the patient BC4. The b- and y-series fragment ions identified in the Sequest database search were displayed in red and blue letters, respectively. The centroid MS/MS spectrum of endogenous TKT_120-132_ peptide detected from urinary EVs (B) or synthetic TKT_120-132_ peptide (C) was shown in the left panel. The right matrix demonstrates the sequence coverage in the Sequest database search. The b- and y-series fragment ions identified in the Sequest database search were displayed in red and blue letters, respectively.

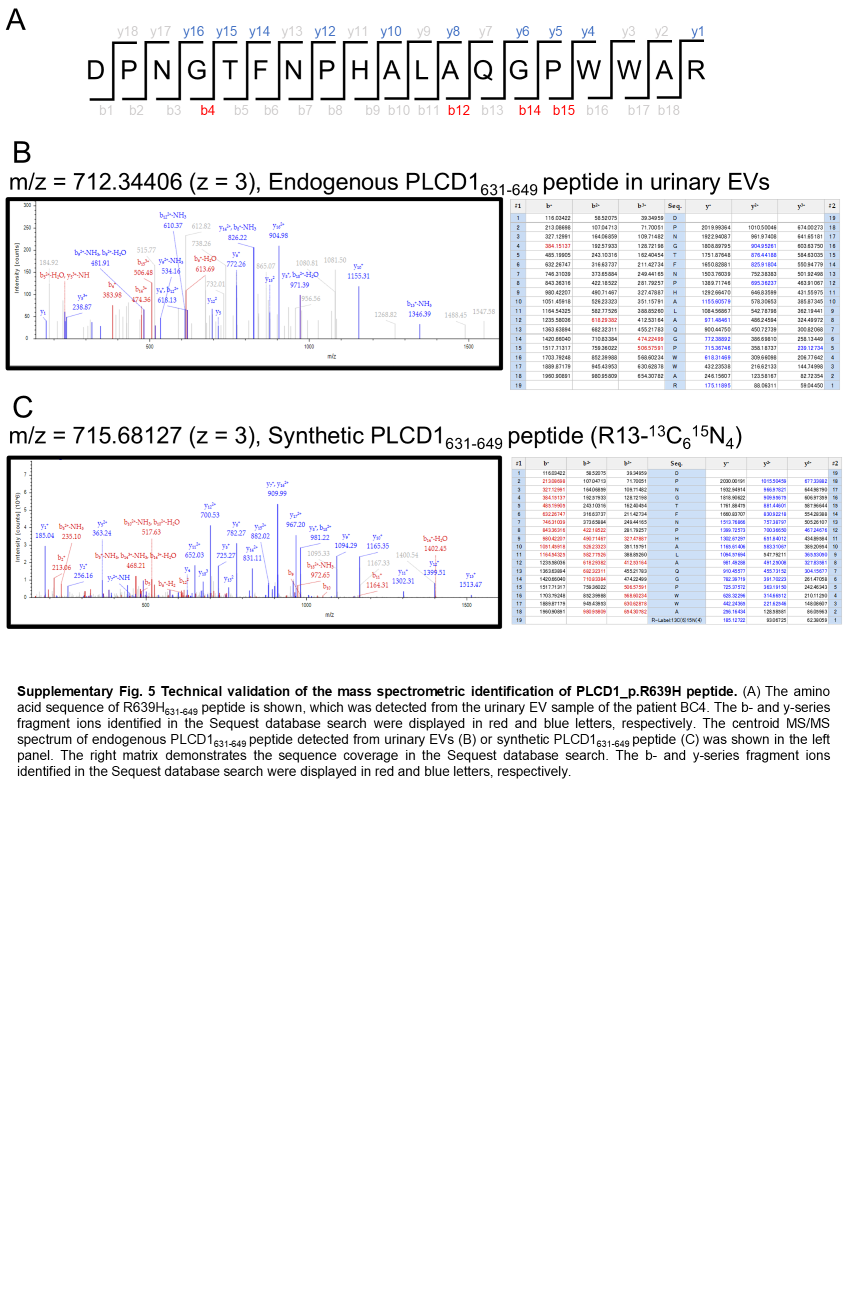
Fig. S5. Technical validation of the mass spectrometric identification of PLCD1_p.R639H peptide. (A) The amino acid sequence of R639H_631-649_ peptide is shown, which was detected from the urinary EV sample of the patient BC4. The b- and y-series fragment ions identified in the Sequest database search were displayed in red and blue letters, respectively. The centroid MS/MS spectrum of endogenous PLCD1_631-649_ peptide detected from urinary EVs (B) or synthetic PLCD1_631-649_ peptide (C) was shown in the left panel. The right matrix demonstrates the sequence coverage in the Sequest database search. The b- and y-series fragment ions identified in the Sequest database search were displayed in red and blue letters, respectively.

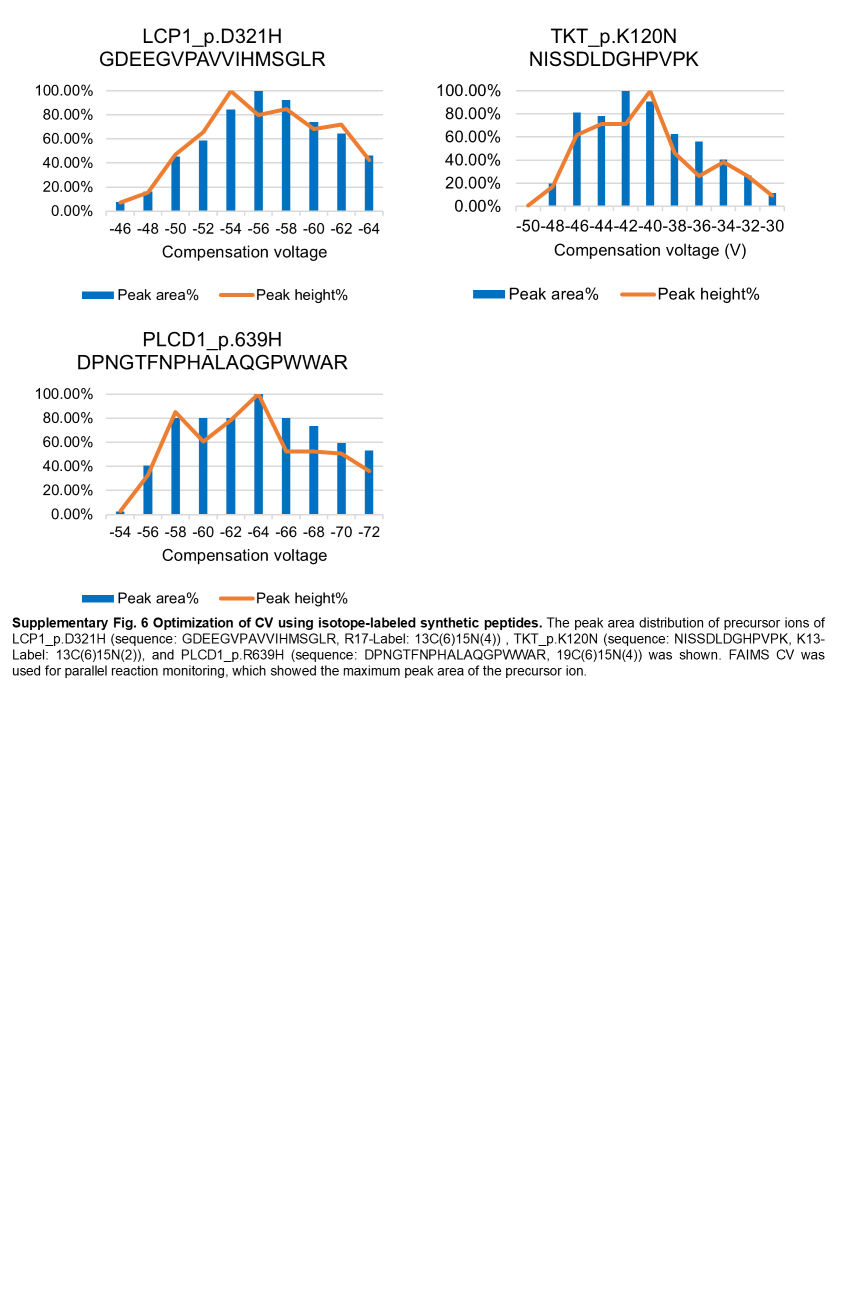

Fig. S6. Optimization of CV using isotope-labeled synthetic peptides. The peak area distribution of precursor ions of LCP1_p.D321H (sequence: GDEEGVPAVVIHMSGLR, R17-Label: 13C(6)15N(4)) , TKT_p.K120N (sequence: NISSDLDGHPVPK, K13-Label: 13C(6)15N(2)), and PLCD1_p.R639H (sequence: DPNGTFNPHALAQGPWWAR, 19C(6)15N(4)) was shown. FAIMS CV was used for parallel reaction monitoring, which showed the maximum peak area of the precursor ion.

Table S1. Clinical characteristics of bladder cancer patients

Table S2. List of identified proteins in bladder cancer tissues

Table S3. List of identified proteins in cultured tissue-derived EVs (CTD-EVs)

Table S4. List of identified proteins in urinary EVs

Table S5. List of identified mutant peptides in cancer tissues

Table S6. List of identified mutant peptides in cultured tissue-derived EVs (CTD-EVs)

Table S7. Parameters of parallel reaction monitoring (PRM) for mutant peptides
